# Large Reaching Datasets Quantify the Impact of Age, Sex/Gender, and Experience on Motor Control

**DOI:** 10.1101/2025.05.01.651757

**Authors:** Aoran Zhang, Marit F. L. Ruitenberg, Matthew Warburton, Stephen Scott, Jonathan S. Tsay

## Abstract

As we age, our movements become slower and less precise—but the extent of this decline remains unclear. To address this, we harmonized data from 2,185 participants across four published studies using a standard center-out reaching task. We found that older age was associated with a steady decline in reaction time (– 1.2 ms/year), movement time (–2.3 ms/year), and movement precision (–0.02º/year). Although the rate of decline did not differ by sex/gender, females consistently reacted more slowly (–8 ms), moved more slowly (–37 ms), and exhibited greater precision (+0.5º) across the adult lifespan. Notably, sex/gender differences attenuated after accounting for experiential factors such as video game use and the amount of sleep per day, whereas age remained a robust and consistent predictor of motor decline. Together, these findings provide a large-scale quantification of age, sex/gender, and experiential effects on motor control, offering a normative benchmark to inform future clinical interventions aimed at preserving motor function across the lifespan.

## Introduction

As we get older, our movements become slower and less precise (Bielak et al., 2014; Der & Deary, 2006; Dominika Dykiert et al., 2012; Fozard et al., 1994; Hultsch et al., 2002; Myerson et al., 2007; Salthouse, 2000; Seidler et al., 2010; Stirling et al., 2013). But exactly how much remains unclear for several reasons. First, many studies on age-related slowing rely on simple reaction time tasks, typically measured by the speed of button presses (Bianco et al., 2020; Deary et al., 2011; Dominika Dykiert et al., 2012; Fozard et al., 1994; Hultsch et al., 2002; Salthouse, 2000; Yuan et al., 2008). Given that these tasks place minimal demands on precision and often conflate distinct components of motor control (e.g., reaction time & movement time), they offer only a narrow view into the impact of aging on motor performance.

Second, many studies on age-related slowing use complex psychophysical tasks that often place substantial cognitive demands. For instance, older adults often move more slowly and less precisely when continuously tracking a moving target (Baweja et al., 2015; Jagacinski et al., 1995; Liutsko et al., 2020; Pratt et al., 1994b; Stirling et al., 2013). However, these tasks tap into more than just motor control—they may require sustained attention, working memory, and the ability to predict the target’s motion (Crundall et al., 2008; Marvel et al., 2019; Mathew et al., 2018; Stuhr et al., 2018; Vaziri et al., 2006). This makes it unclear whether the age-related deficits arise from challenges in *deciding* where to move, or from *executing* the movement itself (Elliott et al., 2010; Miall et al., 1993; Morgan et al., 1994; Pratt et al., 1994a).

Third, studies using simple center-out reaching tasks—a standard method for quantifying motor control— often lack the statistical power to detect an age-related effect. Typically, these studies are conducted in person with highly precise motion-sensing equipment but include fewer than 100 participants (10 individuals/decade); most studies divide people into arbitrary “young” and “old” categories, often with fewer than 20 participants per group, ignoring how age may impact motor performance across the continuous lifespan.

Compounding this issue, most studies have overlooked the contributions of sex/gender and experience on motor performance. Evidence for sex/gender differences is controversial: some studies report faster and more precise movements in men compared to women (Bianco et al., 2020; Der & Deary, 2006; Dorfberger et al., 2009; Dominika Dykiert et al., 2012), whereas other studies find the opposite pattern (Jiménez-Jiménez et al., 2011; Ruitenberg et al., 2023; Stirling et al., 2013; Yuan et al., 2008). Moreover, experiential factors—such as video game use or sleep quality—have not been systematically examined. Neglecting these factors is problematic not only because they may shape the neural and behavioral foundations of motor control (Bianco et al., 2020; Dorfberger et al., 2009; Jiménez-Jiménez et al., 2011; Lissek et al., 2007), but also because their potential interaction with age complicates interpretations of age-related effects (Ghisletta et al., 2018; Roivainen et al., 2021).

To fill this gap, we re-analyzed data from 2,185 participants across four published studies using a standard center-out reaching task—yielding one of the largest datasets of its kind. This large sample, combined with Bayesian multilevel modeling, gave us the statistical power to precisely quantify how aging impacts reaction time, movement time, and spatial precision across the adult lifespan. We treated age as a continuous variable, controlled for sex/gender, and examined the role of experiential factors such as video game use and sleep. Together, these findings offer a large-scale characterization of motor control across age and sex/gender, establishing a normative benchmark to guide future clinical interventions aimed at preserving motor function across the adult lifespan.

## Results

We asked a simple but foundational question: How rapidly does motor control decline with age? To answer this, we conducted a re-analysis of four large-scale studies that met five strict inclusion criteria (see Methods). These studies yielded a dataset of 2,185 participants spanning a wide age range (18 to 90 years old) and a balanced sex/gender distribution (F/M: 0.52/0.48)—making it one of the richest collections of reaching data to date (Table 1).

**Table 1.** Summary of the four datasets included in our reanalysis. Each dataset varied in sample size (N), participant demographics (age, sex/gender), geographic location of data collection, experimental apparatus, and task parameters. Apparatus ranged from robotic manipulanda to consumer input devices (e.g., mouse, joystick, stylus). Key task features—including number of targets, target distance, and average performance metrics (reaction time, movement time, and movement precision measured as the standard deviation of movement angle)—are listed for each study. Values are reported as means [95% range of the sample]. Note that Coderre et al. (2010) was part of a larger dataset from the LIMB lab, led by Dr. Stephen Scott, and obtained via personal communication. The data were collected using a combination of Kinarm exoskeleton and endpoint robots.

| Feature/Study | 1. Coderre et al, 2010 | 2. Tsay et al, 2024a | 3. Ruitenberg et al, 2023 | 4. Taylor et al, 2017 |
| --- | --- | --- | --- | --- |
| N | 538 | 1,227 | 157 | 263 |
| Age | 43.2 [19.4, 80] | 29.6 [18, 74] | 37.2 [19.9, 60.4] | 52.9 [21.3, 82] |
| Sex/Gender (F:M) | 308:230 | 621:606 | 79:78 | 135:128 |
| Location | Canada | USA | The Netherlands | UK |
| Apparatus | Robotic Manipulandum | Trackpad/Mouse | Dual-axis Joystick | Stylus |
| Number of Targets | 4 or 8 | 1 | 8 | 4 |
| Target Distance (cm) | 5 | 6 | 5 | 5 |
| Reaction Time (ms) | 311.4 [236.4, 454.7] | 296.4 [163.3, 476.9] | 509.9 [415.8, 639.7] | 310 [263.6, 388.2] |
| Movement Time (ms) | 1056.7 [797.5, 1364.0] | 200.2 [55.0, 581.6] | 384.2 [230.9, 574.6] | 412.4 [314.3, 534.5] |
| Movement Angle SD (°) | 5.7 [1.3, 21.3] | 3.8 [1.6, 7.3] | 3.8 [2.2, 7.7] | 9.4 [4.8, 22.8] |

We focused on three core indicators of motor control. Reaction time—the delay between target onset and movement initiation—reflects sensorimotor readiness. Movement time—the duration of the movement— indexes the efficiency of motor execution. Movement precision, captured by the standard deviation of movement angle, reflects the consistency/noise of motor output. These measures are uniquely captured by center-out reaching tasks, as simple button presses place minimal demands on sensorimotor precision, whereas complex motor tasks often introduce cognitive confounds such as attention, working memory, and decision-making.

As shown in Table 1, reaction time, movement time, and movement precision vary across studies, likely reflecting differences in equipment (Figure 1A), task protocols (Figure 1B), and participant sampling. Rather than treat this heterogeneity as noise, we leveraged it using Bayesian multilevel regression (Chen et al., 2019; Neil et al., 2025), which allowed us to harmonize data across diverse experimental contexts, enabling us to examine the effects of age and sex/gender while properly accounting for study-specific variability. As a result, it yields more precise and generalizable inferences than any single study could provide.

**Figure 1.**
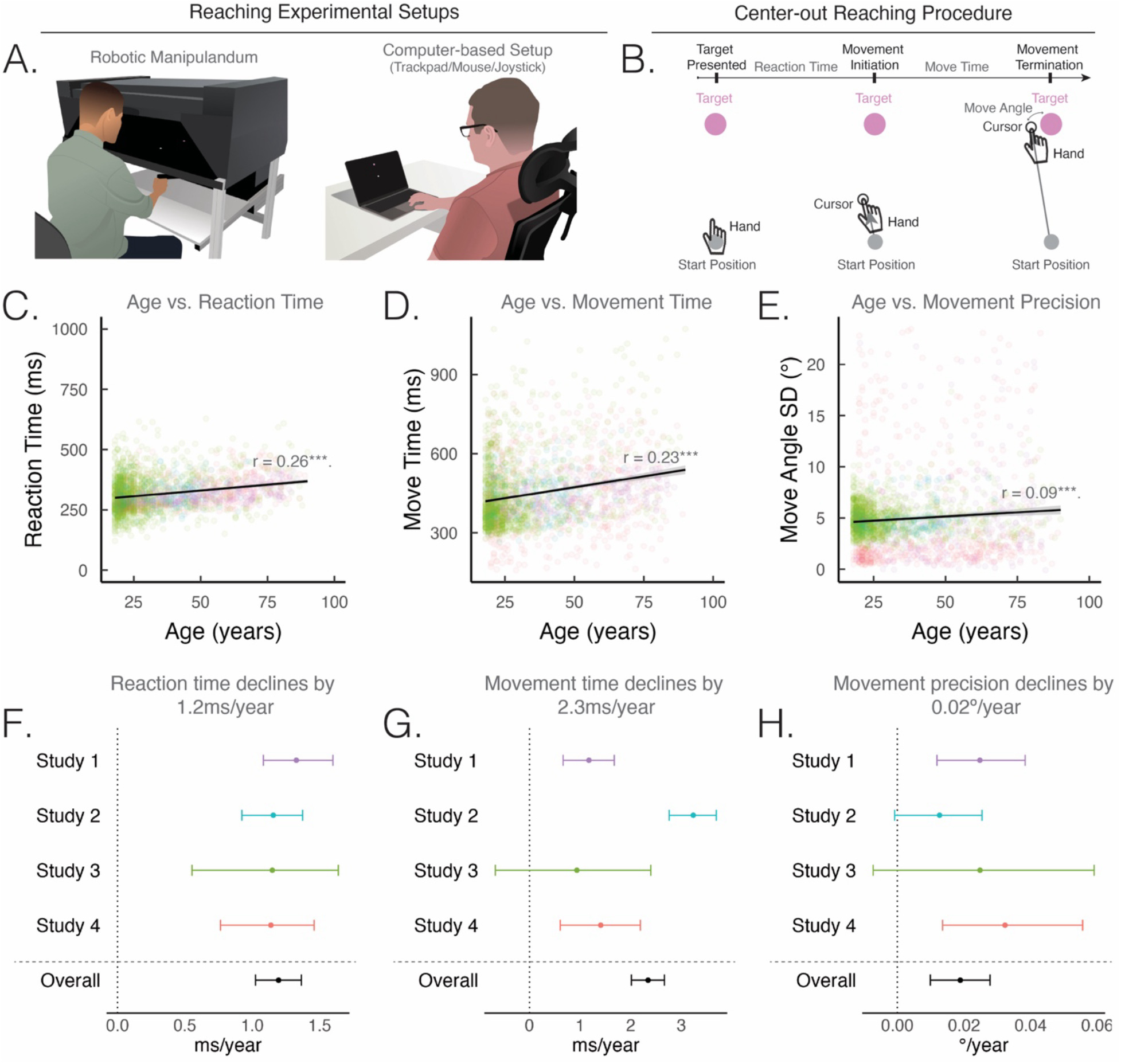
Motor control declines gradually with age. **(A**) Experimental setups from four large-scale reaching datasets. **(B)** In a standard center-out reaching task, a visual target appears (pink circle), prompting participants to initiate a movement. The trial terminates when the visual cursor (black circle) crosses the target distance. Reaction time is defined as the delay from target onset to movement initiation; movement time as the time from movement onset to target crossing; movement angle as the angle of the hand at target crossing; and movement precision as the standard deviation of movement angle across trials. **(C-E)** Reaction time **(C)**, movement time **(D)**, and movement precision **(E)** all exhibit a gradual decline with age. To enhance cross-study comparability and visualization, we normalized each dependent variable by subtracting its study-specific mean and adding the grand mean across all datasets. This approach aligns the scales while preserving meaningful between-group differences. **(F–H)** Bayesian estimates of the effect of aging on motor control. Dot denotes the mean posterior estimate; error bars denote the 95% credible interval of the posterior distribution. Positive values reflect increases in reaction time, movement time, and movement precision, after accounting for study-level differences.

### Aging Degrades Motor Control

It is well-known that motor control slows with age—but our dataset quantifies this decline with unprecedented precision. As shown in Figures 1C–H, reaction time increases by about 1.2 ms per year ([1.0, 1.5]), movement time by 2.3 ms per year ([2.0, 2.7]), and movement precision declines by 0.02º per year ([0.01, 0.03]). (Values reflect posterior means and 95% credible intervals from our Bayesian multi-level model.). These annual changes may seem small, but they compound: Between ages 18 and 90, people become 85 ms slower to react, 170 ms slower to move, and 1.4º less precise—differences that can mean the difference between reaching for a cup or knocking it over, or ability to react appropriately when driving a car.

Despite differences in apparatus, settings, and populations (Table 1), the results were remarkably consistent. All four studies showed the same pattern: reaction and movement times increased with age, while precision declined. Notably, this aging pattern persisted even when the largest dataset (Tsay et al., 2024) was excluded: reaction time increased by about 1.3 ms per year ([1.1, 1.4]), movement time by 1.1 ms per year ([0.7, 1.6]), and precision declined by 0.03º per year ([0.01, 0.05]), underscoring the robustness of these findings.

We also found that performance declined *linearly* across all measures: That is, adding a quadratic term for Age did not improve model fit (ΔBIC (linear – quadratic): Reaction Time = -14.5; Movement Time = -5.5; Movement Precision = -13.4. Negative ΔBIC values indicate a better fit for the linear model; positive values favor the quadratic model.). Taken together, these findings suggest a steady deterioration in motor control with age.

### Males prioritized speed, whereas females favored precision

The impact of sex/gender on motor control is controversial, with some studies reporting no differences and others finding marked disparities (Bianco et al., 2020; Der & Deary, 2006; D. Dykiert et al., 2012; Jiménez-Jiménez et al., 2011; Liutsko et al., 2020; Stirling et al., 2013; Yuan et al., 2008). Leveraging our large, well-powered dataset, we set out to examine these differences with unprecedented precision.

We found that age-related declines in motor performance were similar across sexes/genders. Adding an interaction term between Age and Sex/Gender did not improve model fit (ΔBIC, no interaction – interaction: Reaction Time = -13.2; Movement Time = -18.7; Movement Precision = -15.6), and Bayesian inference confirmed that the Age × Sex/Gender interaction effect was not robust.

However, there were notable Sex/Gender differences in motor performance: Females were 8.0 ms [2.8, 13.3] slower to react and 37.0 ms [26.4, 47.6] slower to move—but were 0.5° [0.2, 0.8] more precise than males (Figures 2D-F). This pattern reveals a classic speed–accuracy trade-off: Males prioritized speed at the expense of precision, while females favored precision over speed.

**Figure 2.**
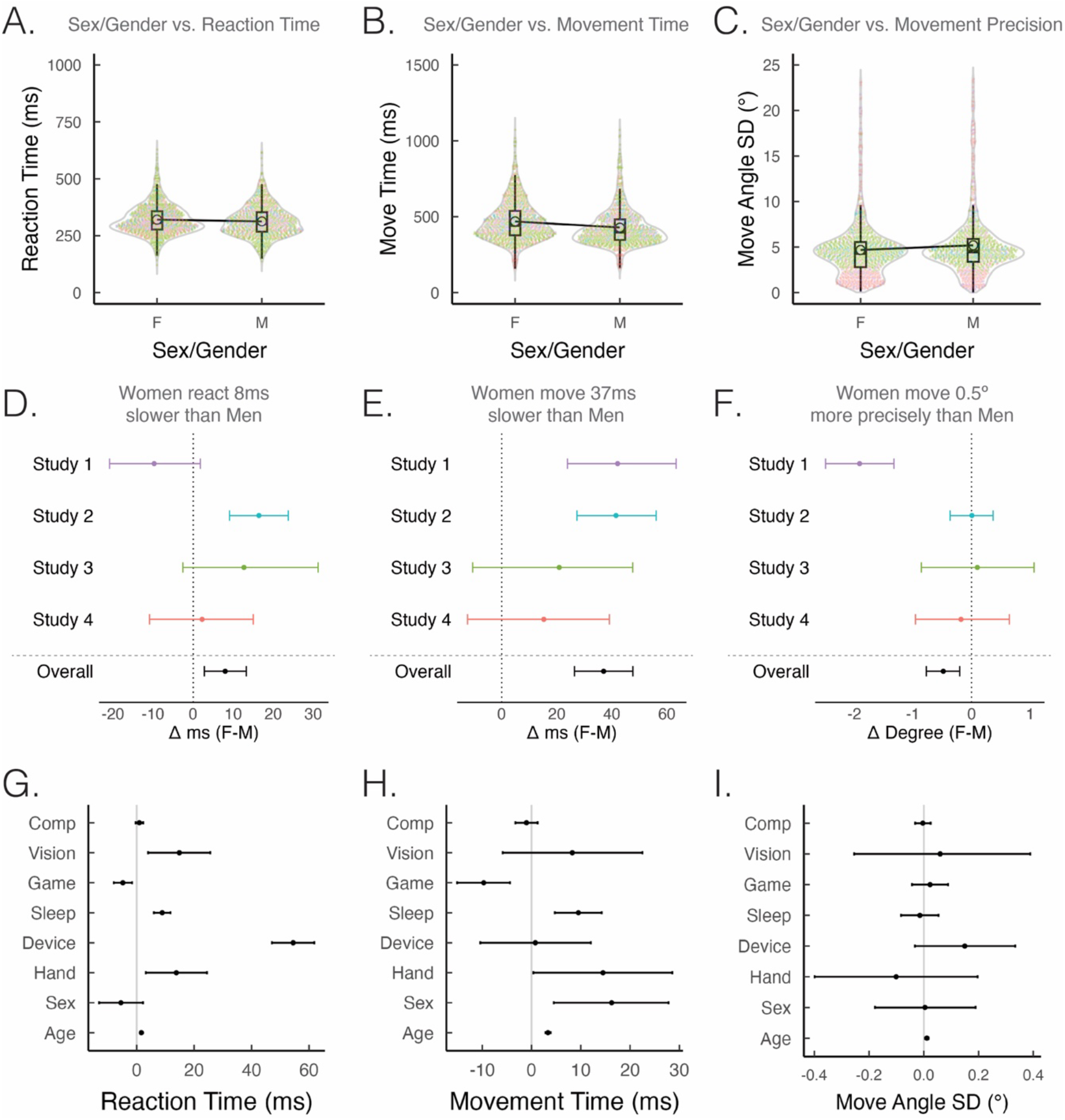
The Impact of Sex/Gender and Experiential Factors on Motor Control. (A–C) Reaction time. **(A)**, movement time **(B)**, and movement precision **(C)** differ across sex/genders. To improve comparability across studies, we normalized each measure by subtracting its study-specific mean and adding the grand mean across all datasets. **(D–F)** Bayesian estimates of sex/gender effects on motor control. Dots indicate posterior means; error bars show 95% credible intervals. Positive values reflect increases in reaction time, movement time, or movement precision, after adjusting for study-level differences. **(G–I)** Exploratory analysis from Tsay et al. (2024) assessing how biological and experiential factors shape motor control. Age effects remained robust across all variables, consistent with a biological basis for motor decline. In contrast, sex/gender effects largely disappeared after accounting for experiential variables such as input device (trackpad vs. mouse), video game experience, vision quality, sleep, and computer usage. See Methods for full variable definitions and modeling details.

### Age-Related Declines Reflect Biology; Sex/Gender Differences Reflect Experience

While age- and sex/gender-related differences in motor control may reflect biological factors, we hypothesized that they may also stem from differences in lived experience. For instance, older adults may favor different input devices (e.g., mouse vs. trackpad), and males and females often differ in video game exposure (Lucas & Sherry, 2004; Rehbein et al., 2016)—both of which can influence motor performance. To test this, we conducted a follow-up analysis of Tsay et al. (2024), a large dataset (N = 1,227) containing detailed self-reported measures of experiential factors, including sleep, computer usage, vision, video game experience, mouse type, and handedness.

Strikingly, age remained a robust predictor of motor performance even after accounting for experiential factors (Figures 2G–I): Reaction time slowed by 1.6 ms/year [1.3, 1.9], movement time by 3.3 ms/year [2.8, 3.9], and movement precision declined by 0.01°/year [0.01, 0.02]. In contrast, sex/gender differences largely disappeared—there was no clear evidence for effects on reaction time (–5.5 ms [–13.1, 2.2]) or movement precision (0.004° [–0.18, 0.19]), as both 95% credible intervals included zero. Although females continued to move slightly more slowly than males (16.3 ms [4.5, 27.8]), this difference was reduced by 55% after adjusting for experiential factors. Together, these findings suggest that age effects on motor control may be more biologically grounded, while sex/gender differences are largely attributable to experiential factors.

We also uncovered several novel experiential influences on motor control (Figures 2G–I). Mouse users responded significantly faster than trackpad users (−54.5 ms [−61.8, −47.1]) and moved slightly more quickly (−0.8 ms [−10.4, 12.0]). Using the dominant-hand, right-handers showed slower reaction times (13.8 ms [3.2, 24.5]) and movement times (14.5 ms [0.37, 28.6]) compared to left-handers. Additionally, each extra hour of sleep was associated with slower reaction (9.0 ms/hour [6.0, 11.8]) and movement times (9.5 ms/hour [4.7, 14.2]). In contrast, greater video game use predicted faster reaction times (−4.8 ms per rating unit [−8.0, −1.6]) and faster movement times (−9.7 ms [−15.1, −4.4]). Together, these findings highlight how everyday experiences can impact motor performance.

## Discussion

Our findings offer one of the most precise quantifications of how motor control changes across the adult lifespan—and how those changes differ by sex/gender and experiential factors. By combining diverse datasets with Bayesian multi-level modeling, we provide a robust and generalizable portrait of motor aging.

### What Drives Age-Related Motor Decline?

We found that reaction time slows by 1.2 ms/year, movement time by 2.3 ms/year, and movement precision declines by 0.02° per year. Motor skills may decline with age for many reasons—and these explanations are not mutually exclusive. One possibility is experiential: In cross-sectional studies like ours, older adults may be less familiar with modern technology, potentially affecting performance. However, even after controlling for experiential factors such as device type, sleep, vision, video game experience, and computer usage, age remained a strong and consistent predictor of slower and less precise movement. This pattern held across datasets and experimental setups, pointing to more fundamental biological drivers of motor decline.

Several biological mechanisms could underlie this decline. On the sensory side, aging may degrade visual acuity and proprioception, making it harder to locate targets or sense limb position. On the neural level, age is associated with reduced white matter integrity (Bennett et al., 2010), disrupted interhemispheric inhibition (Fling & Seidler, 2012), altered functional connectivity (Sala-Llonch et al., 2015), dedifferentiation of neural representations (Cassady et al., 2020), and broader structural and chemical changes in the brain (Seidler et al., 2010). At the periphery, muscle atrophy (Naruse et al., 2023) may impair precision. Even metabolism may play a role: moving quickly becomes more energetically costly with age, making slower movements a rational, energy-conserving strategy (Summerside et al., 2024). Future research could tease apart these influences to pinpoint how much of age-related motor decline reflects experience, how much reflects biology—and why.

### What Drives Sex/Gender Differences in Motor Control?

We found that females tend to react and move more slowly but with greater precision, whereas males react and move more quickly but less precisely. Although the effects on reaction and movement time were consistent across studies, the precision effect was less reliable, highlighting the need for future research to confirm this pattern.

Why might this speed-accuracy trade-off arise? One possibility is that it reflects sex/gender differences in feedback vs feedforward motor control strategies: females may rely more on cautious, online feedback control, while males favor faster, ballistic feedforward control (Bianco et al., 2020). Alternatively, it may reflect a difference in optimal vs robust motor control strategies, with females favoring optimal control (prioritizing accuracy) and males favoring robust control (prioritizing speed). Future work is needed to disentangle these alternatives (Crevecoeur et al., 2019).

Crucially, our results suggest these sex/gender differences may *not* be rooted in biology, but largely in experience. Sex/gender was correlated with experiential variables—such as gaming experience (Festl et al., 2013; Lemmens et al., 2015; Lucas & Sherry, 2004) and device type (Warburton et al., 2024) —and once these were accounted for, the sex/gender effects sharply attenuated. This serves as a caution: apparent sex/gender dimorphisms may in fact reflect experiential ones (Baenninger & Newcombe, 1989, 1995; Helgeson, 2025). Rather than asking whether males and females differ, we should ask why—and look more closely at the roles of experience and context in shaping motor performance.

### What is the impact of experience on motor control?

We found that several experiential factors—such as video game experience, computer usage, and sleep—significantly influenced motor performance. Participants with more video game experience tended to respond and move more quickly and precisely. This may reflect enhanced visuomotor coordination, particularly among those who favor role-playing and shooting games, which demand rapid reactions and fine eye–hand coordination (Bisoglio et al., 2014; Green & Bavelier, 2015; Griffith et al., 1983; Joseph & Willingham, 2000; Listman et al., 2021; Rehbein et al., 2016).

These findings suggest that individual differences in experience are not just statistical noise—they shape sensorimotor behavior in meaningful ways. They should be considered when establishing normative benchmarks for human performance and may serve as a starting point for probing how real-world experiences drive plasticity in sensorimotor systems—and how such changes manifest in the brain. These findings also point to the need for future research to investigate why and how these experiential factors influence motor performance.

### Toward a Normative Benchmark for Sensorimotor Health

We chose the center-out reaching task as a representative measure of motor control because it strikes a balance between simplicity and precision. Unlike complex tasks like trajectory tracking, it minimizes cognitive demands; unlike gross motor tasks such as sit-to-stand and keypress tasks, it enables a fine-grained dissection of distinct phases of motor control. By precisely quantifying how age, sex/gender, and experience influence sensorimotor control, this approach provides a foundation for future studies to revisit more complex tasks and disentangle effects driven by core motor control changes from those reflecting cognitive or other processes.

Although our studies used different input devices—leading to expected differences in mean motor performance (see Table 1)—the effects of aging were strikingly consistent across datasets. To rigorously account for both study-specific and general effects, we applied a Bayesian multilevel modeling (A. Gelman & Hill, 2007; Andrew Gelman et al., 2003; McElreath, 2018) approach to harmonize the data—a method increasingly recognized as best practice in large-scale behavioral research. While input device introduces some variability, the consistent pattern of results across studies underscores the robustness of these effects, suggesting they reflect differences in motor control rather than artifacts of hardware differences.

Despite its size, our dataset does not fully represent the general population. Left-handers are underrepresented, we lack data on non-dominant hand performance, and although socioeconomic status was not measured, our sample likely skews wealthy. We intentionally excluded children to avoid confounding developmental changes with aging effects—possibly explaining why our findings revealed linear rather than quadratic trends. Nonetheless, expanding this ‘living’ dataset to include children and other underrepresented groups is crucial for building a comprehensive normative benchmark of sensorimotor control across all ages, sexes/genders, geographies, and experience levels.

To achieve this, we propose three complementary strategies. First, expand the dataset through coordinated data-sharing initiatives. Second, integrate precision tools—such as the Kinarm robot—into routine clinical assessments, akin to standard measures like blood pressure (Coderre et al., 2010; Fooken et al., 2023; Scott & Dukelow, 2011; Semrau et al., 2018; Tsay, Steadman, et al., 2024). Third, incorporate scalable online motor assessments to enable cost-effective monitoring of sensorimotor health at the population level (Hooyman & Schaefer, 2023; Malone et al., 2023; Roemmich & Bastian, 2018; Saban & Ivry, 2021; Tsay et al., 2021). Together, these efforts can establish a robust normative benchmark for aging, supporting early detection of age-related motor decline and improving diagnosis of movement and cognitive impairments in older adults (Koppelmans et al., 2024; Raje et al., 2025).

## Methods

### Study identification

To examine the impact of aging and sex/gender on motor control, we identified published datasets that met five strict inclusion criteria: (1) The study used an upper-extremity center-out reaching task, a canonical paradigm in motor control research. (2) The study compared performance across age, treating age as a continuous variable to capture gradual changes over the lifespan. (3) The study compared performance across sex/gender, (4) The study included all three key performance measures of motor control—reaction time, movement time, and movement precision (standard deviation of movement angle). These measures did not need to be explicitly reported in the original manuscript but had to be available in the raw data. (5) The study had a sample size of at least 100 participants, ensuring at least 10 individuals per decade for a robust analysis of aging effects. Our search identified four datasets (Table 1; (Tsay, Asmerian, et al., 2024)).

### General Procedure

Each study followed a similar center-out reaching protocol (Figures 1A-B). An example protocol is provided below from (Tsay, Asmerian, et al., 2024): All participants used their own laptop or desktop computer to access the webpage that hosted the experiment (see a demo of the task at https://multiclamp-c2.web.app/). The participants made reaching movements by moving the computer cursor with their mouse or trackpad (Figures 1A-B). The size and position of stimuli were scaled based on each participant’s screen size. For ease of exposition, the stimulus parameters reported below are for a typical monitor size of 13 inches (1,366 × 768 pixels), and the procedure reported below is for the onetarget version of the task.

On each trial, participants executed a planar movement from the center of the workspace to a peripheral target. The start position was marked by a white annulus (0.5 cm diameter), and the target was indicated by a blue circle of the same size, positioned 6 cm away. Each participant was assigned a single target location, randomly selected from eight possible positions (cardinal: 0°, 90°, 180°, 270°; diagonal: 45°, 135°, 225°, 315°) and remained consistent throughout the experiment.

To initiate each trial, the participant moved the cursor, represented by a white dot on their screen, into the start location. Once the participant maintained the cursor in the start position for 500 ms, the target appeared. The participant was instructed to reach to the target using the cursor. If the movement was not completed within 500 ms, the message ‘too slow’ was displayed in red 20-point Times New Roman font at the center of the screen for 750 ms. The visual cursor was always provided as feedback throughout the entire movement. While all four studies employed a similar center-out reaching procedure, they differed in sample size, apparatus, demographics, and geographical location, as summarized in Table 1.

### Data Pre-processing

We preprocessed the data using the following criteria: (1) We excluded samples with missing demographic information. (2) To isolate aging effects and avoid developmental influences, we restricted the sample to individuals aged 18 and older. (3) We excluded outliers, defined as values falling outside the central 95% of the distribution for each dependent variable (i.e., beyond ±1.96 SD from the mean). (4) We excluded individuals with a history of neurological and/or psychiatric disorders. After preprocessing, we retained 538 of 558 samples from Coderre et al. (2010) (part of a larger dataset from the LIMB lab led by Dr. Stephen Scott acquired via personal communication), 1,227 of 2,282 from Tsay et al. (2024), 157 of 385 from Ruitenberg et al. (2023), and 263 of 318 from Taylor et al. (2017), resulting in a final sample of 2,185 unique participants.

### Dependent variables

To examine how aging affects motor control, we analyzed three key variables: reaction time, movement time, and movement precision (quantified as the standard deviation of hand angle across trials). Reaction time was defined as the interval from target appearance to movement initiation.

Movement time spanned from movement initiation to termination. Movement angle was measured as the cursor’s angular deviation from the target at the moment it crossed the target radius.

Study characteristics—such as movement distance and number of targets—can influence overall performance: longer distances typically increase movement times, and more targets can slow reaction times. Variations in movement onset definitions and apparatus further contribute to differences in absolute performance levels across studies. However, our primary interest lies not in these raw values—the study-specific intercepts—but in how performance changes with age, reflected in the slope of these relationships. Since methodological differences are unlikely to systematically vary with age, they should not bias age-related trends. Additionally, our hierarchical Bayesian approach explicitly models study-level variability, allowing us to harmonize data across studies and enhance the robustness and generalizability of our conclusions.

### Bayesian Multilevel Modeling

We used a hierarchical Bayesian statistical model to harmonize our four datasets. Our model includes the following parameters:

‐ Overall (population-level) intercept: *α* ∼ N(0, 10).
‐ Overall Age Effect: *β*_Age_ ∼ N(0, 10).
‐ Overall Sex/Gender Effect: *β*_Sex/Gender_ ∼ N(0, 10).
‐ Residual Overall Error (σ): σ ∼ Ca(0, 2.5).
‐ Study-specific Intercepts: *u_α,j_* ∼ N(0, σ*_α_*).
Study-specific Age Effects: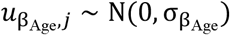.
‐ Study-specific Sex/Gender Effects: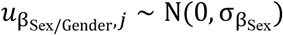.
‐ Residual Study-specific Error: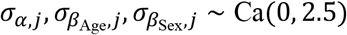.

These priors are chosen to be weakly informative so that the data drive the inference with reasonable constraints (Andrew Gelman, 2006; Andrew Gelman et al., 2008; Polson & Scott, 2012). We tested alternative priors, and the pattern of results remained consistent. *j* here denotes the index of the study each observation is associated with, where *n* is the index of the individual observation. N denotes a normal distribution, and Ca denotes a half Cauchy distribution. With the parameters, our multilevel Bayesian model is defined as follows:

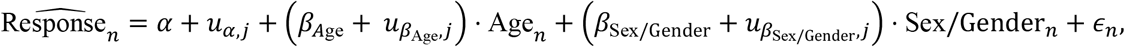

where *ϵ_n_* ∼ N(0, σ) represents the residual error. Age and Sex/Gender are covariates (predictors) while response variables include reaction time, movement time, and standard deviation of movement angle.

### Exploratory Analysis of Tsay et al. (2024)

We conducted a linear Bayesian regression (non-hierarchical) to reanalyze data from Tsay et al (2024), incorporating both demographic and experiential variables. In addition to age and sex, the model included: input device (Mouse/Trackpad: 536/691), handedness (Left/Right: 108/1,119), video game frequency (self-reported Likert scale from –2 to 2, where –2 = strongly disagree and 2 = strongly agree with “I play a lot of video games”; Mean [95% Sample Range]: –0.4 [-2, 2]), daily computer usage (6.8 hours [1, 12]), daily sleep (7.1 hours [5, 10]), and self-reported vision status (Intact/Not Intact: 1137/90). Weakly informative priors were specified as follows:

‐ Intercept: *α* ∼ N(0, 10).
‐ Age Effect: *β*_Age_ ∼ N(0, 10).
‐ Sex/Gender Effect: *β*_Sex_∼ N(0,10).
‐ Handedness Effect: *β*_Hand_ ∼ N(0, 10).
‐ Device Effect: *β*_Device_ ∼ N(0, 10).
‐ Vision Effect: *β*_Vision_ ∼ N(0, 10).
‐ Video Games Frequency Effect: *β*_Game_ ∼ N(0, 10).
‐ Sleep Effect: *β*_Sleep_∼ N(0, 10).
‐ Computer Usage Effect: *β*_Comp_∼ N(0, 10).
‐ Residual Error: σ ∼ Ca(0, 2.5).

With these parameters, our Bayesian model is defined as follows:

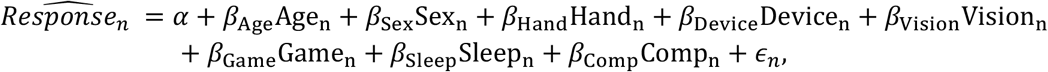

### General Model Fitting Procedure

We implemented our models in Rstan (Carpenter et al., 2017). Stan uses the No-U-Turn Sampler (NUTS), an adaptive variant of Hamiltonian Monte Carlo (HMC) that efficiently explores high-dimensional parameter spaces while avoiding the random walk behavior of traditional MCMC methods (Hoffman & Gelman, 2011). For Bayesian inference, we ran 8,000 iterations for each of the four independent chains, discarding the first 4,000 iterations per chain as warm-up to ensure convergence.

